# Grifola frondosa polysaccharide Ameliorates Inflammation and Insulin Resistance by Regulating Macrophage Polarization of liver in Type 2 diabetes mellitus Rats

**DOI:** 10.1101/2024.05.21.595247

**Authors:** Pei Zou, Xueyan Li, Liping Wang, Ying She, Chenyang Xiao, Yang Peng, Xu Qian, Peng Luo, Shaofeng Wei

## Abstract

Type 2 diabetes mellitus (T2DM) is a common metabolic disease characterized by a lack of insulin secretion, insulin resistance (IR), and hyperglycemia. Given its high prevalence and multifarious complications, diabetes is globally ranked as the third leading cause of mortality. It is critical to discover efficient medication substitutes in order to lessen the drawbacks and adverse effects of traditional diabetes medication therapy. Previous studies have shown that Grifola frondosa polysaccharide (GFP) have the positive effects in regulating blood glucose and insulin resistance, but the understanding of its regulatory mechanism is still limited. Therefore, this study aimed to investigate the effects of GFP on liver inflammation induced by a high-fat diet (HFD) combined with streptozotocin (STZ) in type 2 diabetic rats and to explore its possible mechanisms. The results showed that GFP intervention reduced weight loss and hyperglycemia symptoms, as well as lowered FINS, HOMA-IR, IPGTT-AUC, and IPITT-AUC to different degrees in T2DM rats. At the same time, after GFP intervention, the secretion level of inflammatory factors (TNF-α, IL-1β, MCP-1) was down-regulated and the secretion level of anti-inflammatory factor (IL-10) was up-regulated in the liver tissue of T2DM rats. Furthermore, GFP reduced macrophage infiltration in liver tissue, inhibited macrophage M1-type polarization, and promoted M2-type polarization. These results suggest that GFP intervention could attenuate the hepatic inflammatory response in T2DM rats; possible mechanisms for this effect include hepatic macrophage infiltration and M1/M2 polarization.

**Summary statement:** This study revealed the improvement effect of GFP on hepatic inflammation and insulin resistance in T2DM rats and to explore the possible key roles of hepatic macrophages and their potential mechanisms.

## Introduction

Type 2 diabetes mellitus (T2DM) is a prevalent chronic metabolic disorder typified by deficiencies in insulin secretion, insulin resistance (IR), and hyperglycemia, thus presenting a substantial threat to human health (Kahn et al., 2014). A growing body of evidence suggests that insulin resistance plays a vital role in the pathogenesis of T2DM. As a critical target organ of insulin, the liver plays a crucial role in controlling blood glucose homeostasis and regulating insulin secretion (Tanase DM et al., 2020). It has been found that when the sustained hyperglycemic state exceeds the metabolic capacity of the liver, the role of the liver in regulating blood glucose becomes imbalanced, which is a crucial factor in the occurrence and progression of T2DM (Agrawal R et al., 2021). Therefore, preserving or protecting liver function is essential for the early prevention and treatment of diabetes. Currently, several studies have discovered that the edible fungus can attenuate liver injury, regulate blood glucose, and improve insulin resistance (Ahmad MF et al., 2023) (Rehman AU et al., 2022), which may be used as a precise nutritional intervention for T2DM.

Currently, the most often used hypoglycemic medications include metformin, acarbose, sulfonylureas, and thiazolidinediones. Nevertheless, prolonged administration of these medications is associated with adverse toxicity and side effects on health (Zheng Y et al., 2017). Therefore, there is an urgent need to find secure and efficacious medication substitutes. Grifola frondosa is an edible and medicinal fungus. The fungal biomass suggests that the substrate possesses high nutritional value, yet it also exhibits a diverse array of bioactivities that have been empirically demonstrated (Wu et al., 2021), including immunomodulatory, antiviral, antidiabetic, anti-inflammatory, and intestinal flora regulation (Li L et al., 2019). In contrast, Grifola frondosa polysaccharides (GFP) are its most abundant and significant active component. Several studies have reported that GFP can improve dyslipidemia, hypertension, and pulmonary fibrosis by improving inflammatory symptoms (Li C et al., 2019). There have been some experimental studies on GFP for antidiabetic and hepatic insulin resistance (Xiao C et al., 2015) (Guo WL et al., 2020), the mechanisms are mainly focused on exploring insulin signaling pathway abnormalities, oxidative stress, endoplasmic reticulum stress, and mitochondrial dysfunction (Zhao H et al., 2022) (Arroyave-Ospina JC et al., 2021) (Hao Y et al., 2021), however, little has been reported on the anti-inflammatory effects and its deep-rooted mechanisms for ameliorating T2DM.

It has been discovered in recent years that macrophages are crucial for the development of T2DM or IR. Several studies have pointed out that abnormal recruitment and polarization of macrophages in tissues play a vital role in the pathogenesis of insulin resistance and T2DM (Xu L et al., 2018); A study revealed that inhibition of the NF-κB pathway influenced macrophage polarization and mitigated T2DM-associated periodontitis (Xia S et al., 2024); and another study showed that activation of PPARγ promoted M2 macrophage polarization, which could attenuate diabetic mice obesity-associated adipose tissue inflammation and insulin resistance (Xu J et al., 2023); In addition, studies have reported that the regulation of hepatic inflammation and lipid metabolism abnormalities in T2DM mice may be influenced by macrophage M1/M2 polarization and mitochondrial dynamics (Wang Y et al., 2022). However, it remains unknown whether GFP influences hepatic inflammation and insulin resistance by modulating macrophage polarization, thereby potentially ameliorating the progression of T2DM. To establish a robust scientific basis for exploring the potential efficacy of phytochemicals in T2DM prevention and treatment, this study attempts to define the improvement effect of GFP on hepatic inflammation and insulin resistance in T2DM rats as well as to investigate the potential critical roles and mechanisms of hepatic macrophages.

## Materials and methods

### Materials

High-fat feed (42.4% basic feed, 15% sucrose, 10% lard, 2.5% cholesterol, 0.1% bile salt, 10% egg, and 20% sesame oil) was purchased from Beijing Biotech Co., Ltd (Beijing, China). Grifola frondosa polysaccharides (GFP) were prepared according to the previous experimental conditions of our research group (Liu X et al., 2023), with a polysaccharide content of 92.50% and β-glucan content of 21.03%. Streptozotocin (STZ, Sigma, USA). Metformin hydrochloride tablets (Shanghai Shiguibao Pharmaceutical Co., Ltd, China). ELISA kits for rat FINS, IL-1β, IL-10, MCP-1, and TNF-α were purchased from Jianglai Biotechnology Co., Ltd (Shanghai, China). Type IV collagenase (Sigma, USA). Rat GM-CSF (PeproTech, USA). Lipopolysaccharides (LPS, Solarbio, China). FITC Mouse Anti Rat CD68 (20060, Santa Cruz Biotechnology, China). APC Mouse Anti Rat CD11b (562102, BD Biosciences, USA). BB700 Mouse Anti Rat CD86 (746002, BD Biosciences, USA). PE Mouse Anti Rat CD206 (58986, Santa Cruz Biotechnology, China).

### Animals

Male SD rats (200 ± 20 g) were purchased from the Experimental Animal Center of Guizhou Medical University, China. All rats were housed in a 12 h alternating light/dark environment, with a temperature of 22 ± 2 ℃ and humidity of 50 ± 5 %; the rats could freely access food and water. All animal experiments were approved by the Experimental Animal Ethics Committee of the Guizhou Medical University of China (Approval No. 202303090).

### Establishment of T2DM model and measurements

After 1 week of adaptive feeding, the rats were randomly divided into the Normal control group (NC, n=12) and High-fat diet group (HFD) and respectively fed with normal or high-fat feed. After 8 weeks, the rats in the HFD group were injected intraperitoneally with 1% Streptozotocin (30 mg kg^-1^) (Skovsø S et al., 2014), and the NC group was injected intraperitoneally with an equal amount of buffer. After 72h following STZ injection, the rats showing fasting blood glucose higher than 16.7 mmol L^-1^ were considered successful T2DM models. Subsequently, the T2DM rats were randomly divided into 3 groups: (1) T2DM model group (DM, n=12), fed high-fat chow and given saline intervention for 8 weeks; (2) Metformin intervention group (MET, n=12), fed high-fat chow and given metformin (200 mg kg^-1^) for 8 weeks; (3) GFP intervention group (GFP, n=12), fed high-fat chow and given GFP (1.5 g kg^-1^) for 8 weeks (**Fig. 1**). The GFP intervention dose was converted to the recommended intake of β-glucan of 3 g per day as recommended by the FDA, according to the β-glucan content of the GFP samples and the equivalent dose relationship between rats and humans. The metformin intervention dose was converted according to the equivalent dose relationship between rats and humans based on the principles of clinical administration in adults (optimal effective dose of 2g per day).

**Fig. 1.**
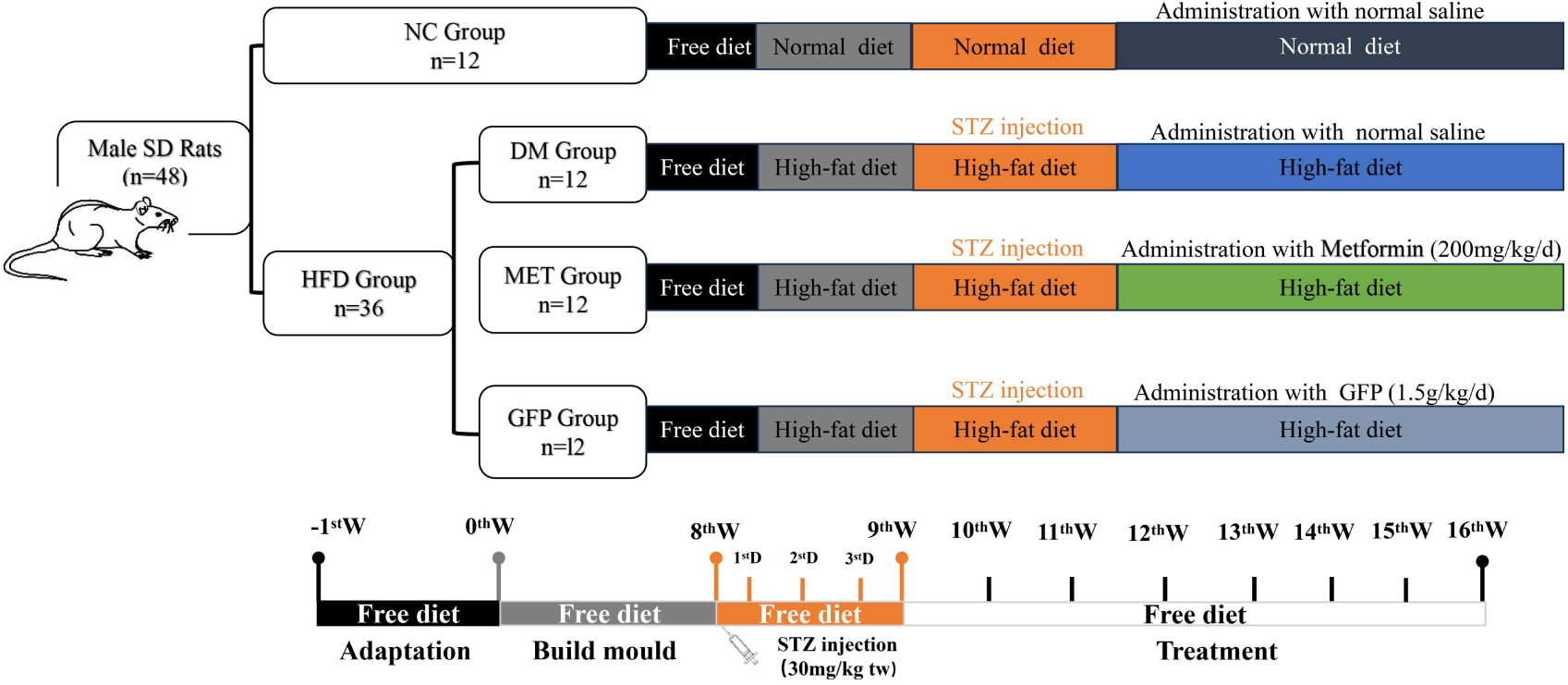
Animal experiment design.

### General symptom monitoring and Fasting blood glucose (FBG) measurement

From the beginning of modeling to the end of the intervention, the rats were closely monitored daily for dietary intake, water intake, hair condition, and mental condition. During the intervention period, each rat’s body weight and FBG were measured once a week.

### Intraperitoneal glucose tolerance test (IPGTT) and Intraperitoneal insulin tolerance test (IPITT)

At the end of the intervention, all rats were fasted with water for 12 h before performing the Intraperitoneal glucose tolerance test (IPGTT), according to the reference method (Bernardi S et al., 2012), all rats were injected intraperitoneally with 50% glucose solution (2 g kg^-1^). For the Intraperitoneal insulation tolerance test (IPITT), the rats were fasted with water for 4 h and then injected intraperitoneally with insulin solution (1 U kg^-1^). Subsequently, the blood glucose levels in the IPGTT and IPITT were measured at 0, 30, 60, 90, and 120 minutes. The areas under the curve (AUC) of IPGTT and IPITT were calculated by GraphPad Prism.

### Assessment of insulin and HOMA-IR

Blood samples were centrifuged to obtain serum, after which the fasting insulin (FINS) was measured using an ELISA reagent kit. The insulin resistance index (HOMA-IR) was calculated according to the following formula: HOMA-IR = [FBG (mmol L^-1^) × FINS (µIU mL^-1^)] / 22.5.

### Histopathological and immunohistochemical analysis

The Liver tissues of rats were fixed in 4% paraformaldehyde, dehydrated, and embedded in paraffin. Subsequently, the liver tissue slides were stained with H&E. All processes were performed according to standard protocols. For immunohistochemistry (IHC), paraffin sections of liver tissue were supplemented with primary antibody (CD68) overnight at 4 ℃, after which the secondary antibody was added for incubation at 37 ℃ for 30 min. Staining was performed using DBA and hematoxylin. Finally, the image was collected by microscope. Macrophage counting was performed by reference method (Pervin M et al., 2021), and five areas were randomly selected from the central vein, portal vein, periportal, perivenular, or Glisson’s sheath of the liver, and cells expressing CD68 with clear nuclei were counted.

### Enzyme-linked immunosorbent assay (ELISA)

An appropriate amount of liver tissue was taken and PBS was added to prepare the tissue suspension, the supernatant was centrifuged and the protein concentration was determined according to the BCA protein concentration determination kit instructions. According to the results of the total protein measurement, an appropriate volume of saline was added to standardize the protein concentration. Sample IL-1β, IL-10, TNF-α, and MCP-1 concentrations were determined according to the ELISA kit instructions and expressed as pg mg^-1^ protein.

### Extraction and identification of liver macrophages

The isolation of liver macrophages was based on a two-step perfusion procedure followed by density gradient centrifugation as previously described (Zhang Q et al., 2023). Liver macrophages were separated from other non-parenchymal cells by selective adhesion in culture medium coated with rat tail collagen for 2 hours. Afterward, CD68 antibody staining was used for purity measurement of liver macrophages by immunofluorescence and flow cytometry.

### Extraction, induced culture, and treatment of BMDM

Extraction of bone marrow-derived macrophages (BMDM) from rats by the reference method (Gorshkova EA et al., 2023), the cells were differentiated in complete medium containing M-CSF (20 ng mL^-1^) for 7 days. On day 6, BMDM was polarized towards the M1 type by lipopolysaccharide (LPS). According to the experimental design, the extracted cells were divided into 4 groups: (1) Normal control group (NC), treated with PBS; (2) Model group (LPS), treated with LPS (100 ng mL^-1^) (Wang S et al., 2021);(3) Low dose of GFP intervention group (LPS+GFP-L), treated with GFP (200 ug mL^-1^) after LPS stimulation; (4) High dose of GFP intervention group (LPS+GFP-H), treated with GFP (400 ug mL^-1^) after LPS stimulation.

### Cell viability test

BMDM were seeded in 96-well plates at a density of 5×10^5^ cells mL^-1^. After adherence, cells were treated with corresponding concentrations of GFP according to the experimental groups, respectively, with 5 replicate wells in each group. After 24 h of incubation, 10 μL CCK-8 solution was added, and incubation was continued for 1 h. The optical density values were measured at 450 nm, and cell viability was calculated.

### Flow cytometry

Hepatic macrophages or BMDM cells were collected by pancreatin, cells were resuspended in FACS buffer, then set up negative control tubes, single label tubes, FMO control tubes, and sample tubes. Each tube was blocked with Fc receptor blocker for 10 min at room temperature, FITC-CD68, APC-CD11b, BB700-CD86, and PE-CD206 flow antibodies were added as required for 30 minutes, and the cells were resuspended in PBS and detected using a BD FACS Canto Ⅱ flow cytometer. The voltage was adjusted according to each single-label tube, the circle gate was set according to each FMO control tube, and the data were collected by flow cytometry and finally analyzed by Flowsjo software.

### Statistical analysis

All data were analyzed using the SPSS 17.0 statistical software and expressed as the mean ± standard error (mean ± SEM).). If the data obeyed normal distribution and satisfied the variance alignment, one-way ANOVA was used for comparison between multiple groups, and the LSD method was used for the pairwise comparison between the groups; if the variance was not aligned, the Dunnett’s t-test was used for the comparison, *P* < 0.05 was considered statistically significant.

## Results

### GFP improved weight loss, hyperglycemia, glucose intolerance, and insulin resistanc in T2DM rats

After 8 weeks of HFD intervention, the rats in the T2DM model group showed a significant decrease in body weight and a significant increase in FBG compared to the NC group (*P* < 0.01). In contrast, the combination of metformin or GFP intervention resulted in a significant increase in body weight and a significant decrease in FBG compared to the model group(*P* < 0.05); there was no significant difference in the effects between the two intervention groups (**Figs. 2A, 2B**).

**Fig. 2.**
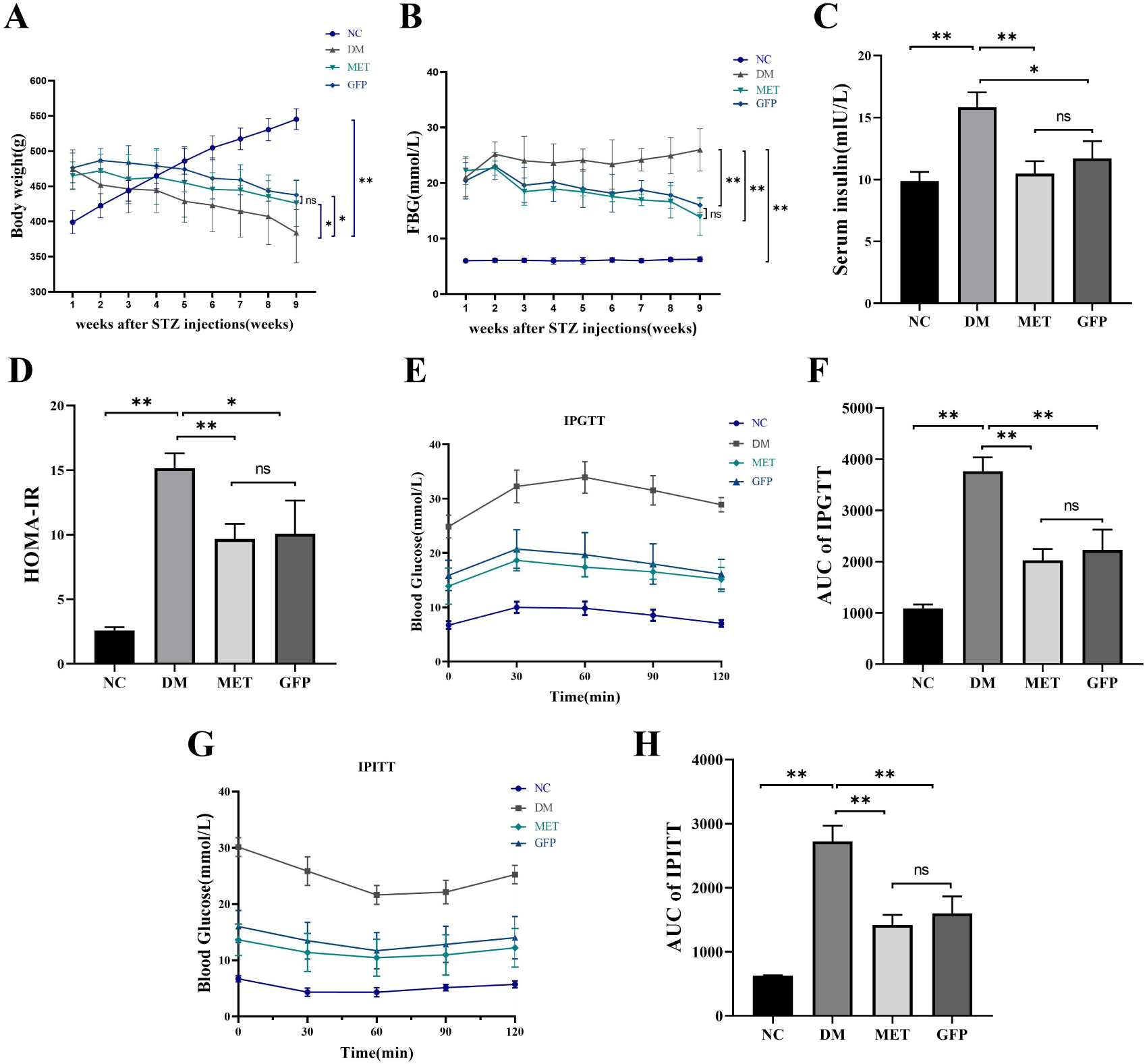
GFP improved weight loss, hyperglycemia, glucose intolerance, and insulin resistance in T2DM rats. (A) The weight changes of rats. (B) The serum fasting blood glucose levels. (C) The serum fasting insulin levels. (D) The insulin resistance coefficient HOMA-IR. (E) Blood glucose levels of intraperitoneal glucose tolerance test (IPGTT). (F) Area under the curve (AUC) of IPGTT. (G) Blood glucose levels of intraperitoneal insulin tolerance test (IPITT). (H) Area under the curve (AUC) of IPITT. The data were presented as mean ± SEM (n=6 in each group). * *P* > 0.05; ** *P* > 0.01; ^ns^ *P* > 0.05.

Compared with the NC group, the FINS, HOMA-IR, IPGTT-AUC, and IPITT-AUC were significantly increased in the model group (*P* < 0.01); Compared with the model group, combining the intervention with metformin or GFP could significantly reduce the FINS, HOMA-IR, IPGTT-AUC, and IPITT-AUC (*P* < 0.05), effectively improved insulin sensitivity in T2DM rats(**Figs. 2C, 2D, 2E, 2F, 1G, 1H**).

### GFP improved hepatic inflammation in T2DM rats

The HE staining results showed that the hepatocytes of rats in the NC group were morphologically intact and neatly arranged, with no obvious degeneration and necrosis, the hepatocyte cords were radially arranged, and the tissue structure was clear, with no obvious inflammatory cell infiltration. In the model group, the arrangement of hepatocytes was disorganized; some hepatocyte nuclei were displaced to the periphery, showing obvious infiltration of inflammatory cells and focal necrosis. The metformin group and the GFP group showed different degrees of improvement compared to the model group (**Fig. 3A**)

**Fig. 3.**
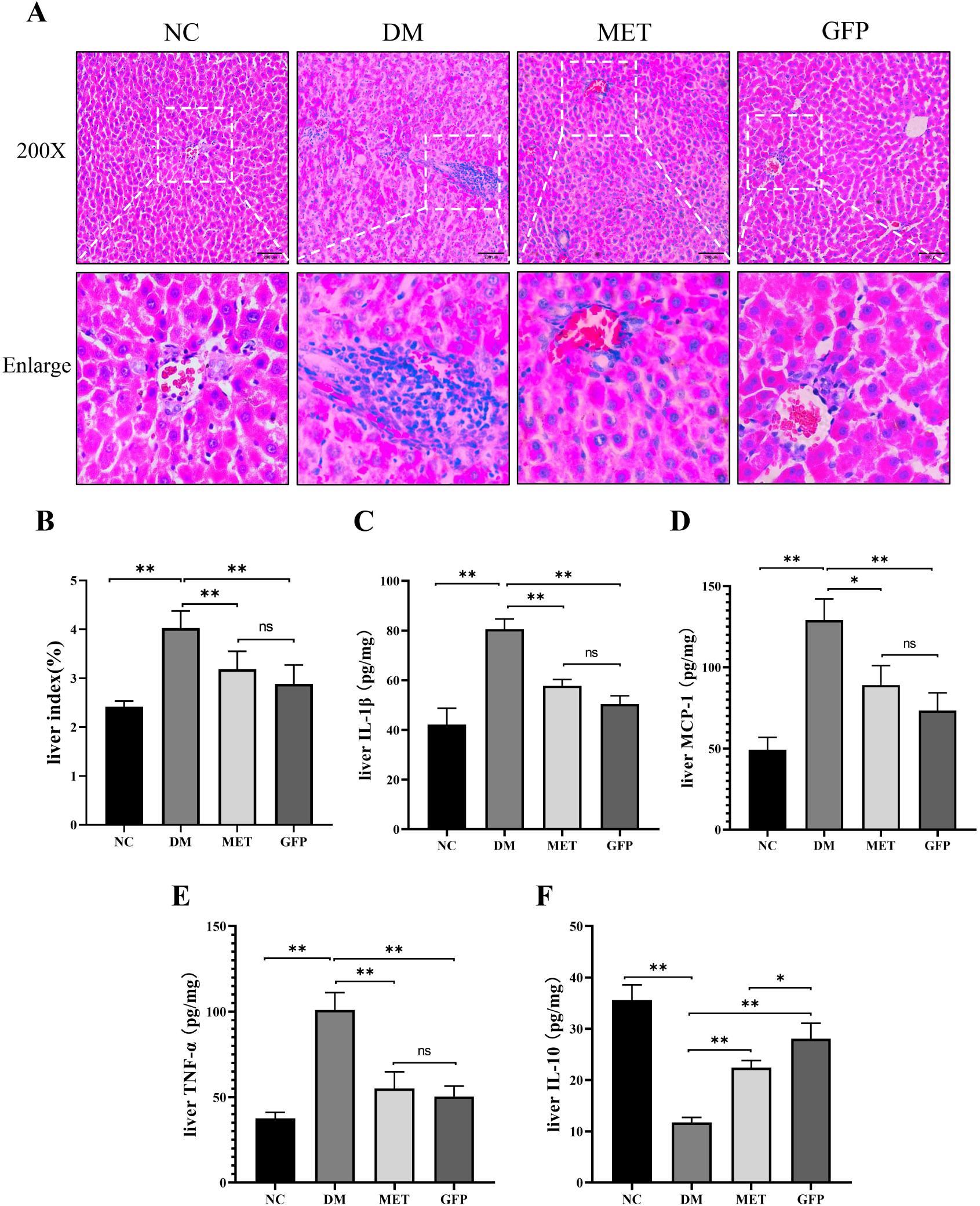
GFP improved hepatic inflammation in T2DM rats. (A) Pathological changes of liver tissue. (B-F) Levels of inflammatory factors in liver tissue. The data were presented as mean ± SEM (n=6 in each group). * *P* > 0.05; ** *P* > 0.01; ^ns^ *P* > 0.05.

Compared with the NC group, the levels of IL-1β, TNF-α, and MCP-1 in the liver tissue of T2DM rats were significantly increased, and the level of IL-10 was significantly decreased (*P* < 0.01). After the intervention of the metformin or GFP, the levels of IL-1β, TNF-α, and MCP-1 were significantly decreased (*P* < 0.05), while the level of IL-10 was increased (*P* < 0.05), and the degree of up-regulation of IL-10 secretion in GFP was significantly higher than that of the metformin group (*P* < 0.05) (**Figs. 3B, 3C, 3D, 3E, 3F**). It was suggested that GFP could improve liver inflammation in T2DM rats.

### GFP decreased macrophage infiltration in the liver tissue of T2DM rats

The immunohistochemical results showed that the 8 weeks of HFD intervention aggravated the infiltration of liver macrophages in T2DM rats compared to the NC group (*P* < 0.05), while the combination of the metformin or GFP intervention could effectively reduce the infiltration of macrophages in T2DM rats (*P* < 0.01), there was no significant difference between the two intervention groups (**Fig. 4**). It was suggested that GFP could effectively improve the degree of macrophage infiltration in liver tissue of T2DM rats.

**Fig. 4.**
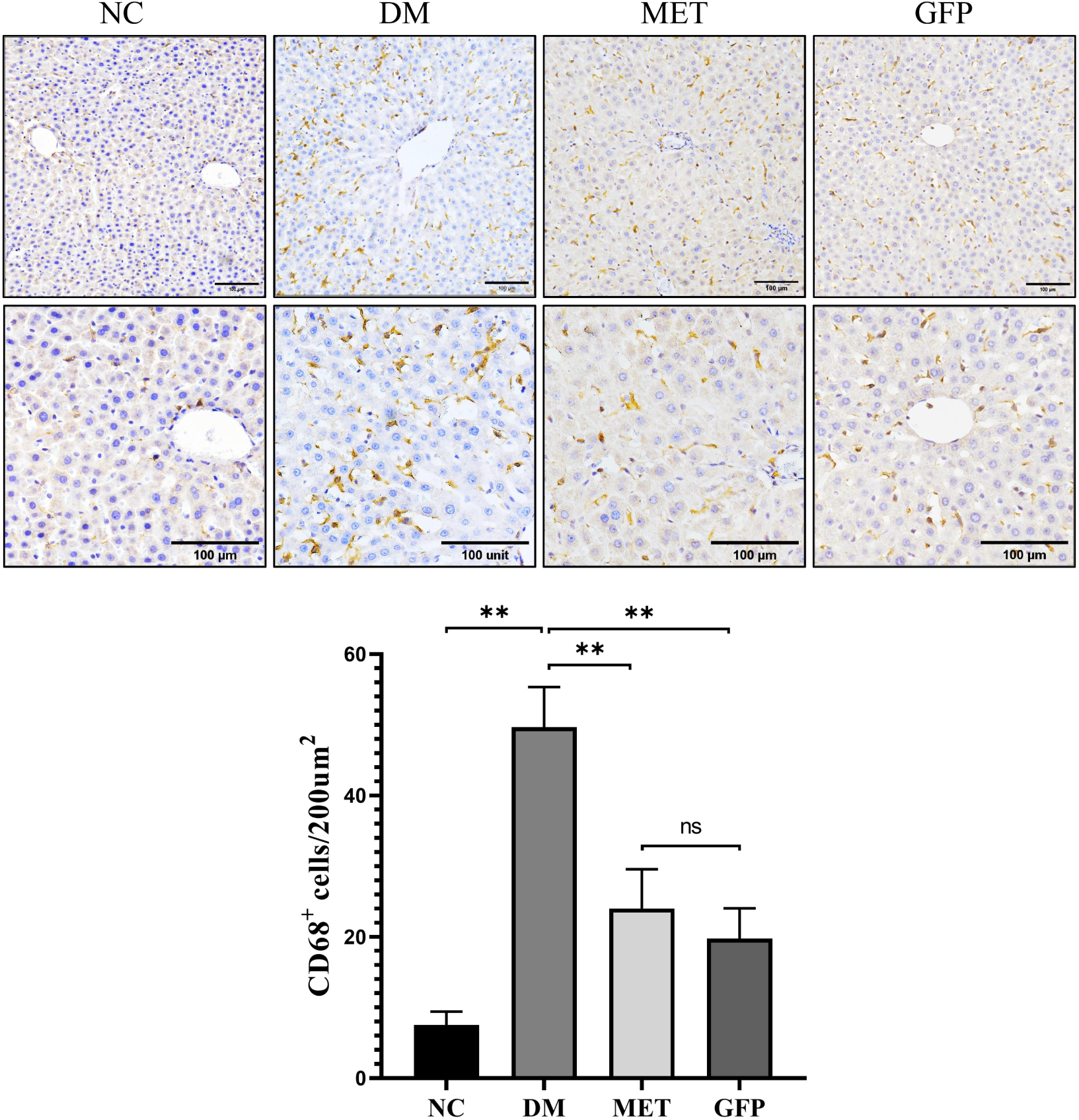
GFP decreased macrophage infiltration in the liver tissue of T2DM rats. The data were presented as mean ± SEM (n=6 in each group). * *P* > 0.05; ** *P* > 0.01; ^ns^ *P* > 0.05.

### GFP inhibited M1-type polarization and promotes M2-type polarization of liver macrophages in T2DM rats

The results of immunofluorescence showed that CD68 was expressed on the membrane of macrophage cells, and the proportion of CD68-positive cells was relatively high (**Fig. 5A**). The results of flow cytometry showed that the proportion of macrophages in the isolated cells was 84.1 ± 3.2 % (**Fig. 5B**), indicated that macrophage isolation was successful, and could be used for the subsequent experiments.

**Fig. 5.**
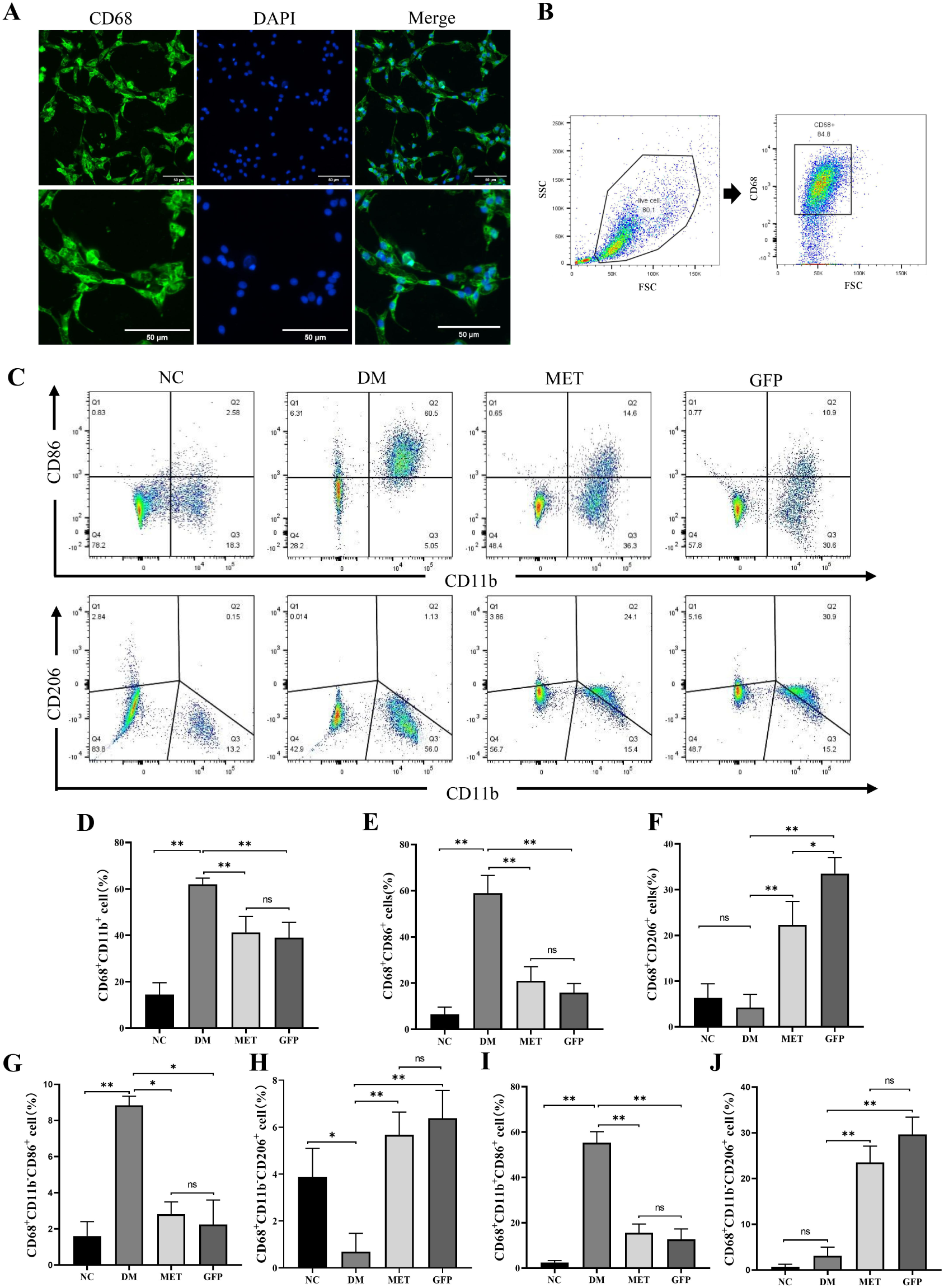
GFP inhibited M1-type polarization and promotes M2-type polarization of liver macrophages in T2DM rats. (A-B) Expression of CD68 was evaluated by immunofluorescence and Flow cytometry in liver macrophages. (C) Flow cytometry analysis of hepatic macrophagel phenotype. (D) The proportion of extrahepatic macrophages. (E) The proportion of overall M1 macrophages. (F) The proportion of overall M2 macrophages. (G)The proportion of intrahepatic M1 macrophages. (H) The proportion of intrahepatic M2 macrophages. (I) The proportion of extrahepatic M1 macrophages. (J) The proportion of extrahepatic M2 macrophages. The data were presented as mean ± SEM (n=6 in each group). * *P* > 0.05; ** *P* > 0.01; ^ns^ *P* > 0.05.

The results of flow cytometry showed that the proportion of peripheral macrophages (CD68^+^CD11b^+^) in the liver tissue of the model group was significantly increased compared with the NC group (*P* < 0.01); after 8 weeks of GFP or metformin intervention, the proportion of peripheral macrophages was significantly increased compared with the model group (*P* < 0.01), there was no significant difference between the two intervention groups (**Figs. 5C, 5D**). The overall phenotype results of liver macrophages showed that the proportion of M1 macrophages (CD68^+^CD86^+^) of the model group was significantly increased compared with the NC group (*P* < 0.01); After 8 weeks of GFP or metformin intervention, the proportion of M1 macrophages was significantly decreased (*P* < 0.01), the proportion of M2 macrophages was significantly increased (*P* < 0.01), and the proportion of M2 macrophages in the GFP group was higher than that in the metformin group (*P* < 0.01) (**Figs. 5C, 5E, 5F**). It was suggested that GFP can achieve anti-inflammatory effects by inhibiting macrophage M1 polarization and promoting M2 polarization.

The phenotype analysis of hepatic intrinsic macrophages showed that the proportion of M1 macrophages (CD68^+^CD11b^-^CD86^+^) was significantly increased (*P* < 0.01) and the proportion of M2 macrophages (CD68^+^CD11b^-^CD206^+^) significantly decreased (*P* < 0.05) in the model group compared to the NC group, after 8 weeks of GFP or metformin intervention, the proportion of M1 macrophages significantly decreased (*P* < 0.05), while the proportion of M2 macrophages significantly increased (*P* < 0.01) (**Figs. 5C, 5G, 5H**). It was suggested that GFP intervention could effectively inhibit the M1 polarization and promote the M2 polarization of liver intrinsic macrophages in T2DM rats.

The phenotype analysis of extrahepatic-derived macrophages showed that the proportion of M1 type macrophages (CD68^+^CD11b^+^CD86^+^) was significantly increased (*P* < 0.01) in the model group of rats compared to the NC group. After 8 weeks of GFP or metformin intervention, the proportion of M1 macrophages was significantly reduced (*P* < 0.01), while the proportion of M2 macrophages (CD68^+^CD11b^+^CD206^+^) was significantly increased (*P* < 0.01) (**Figs. 5C, 5I, 5J**). The above results suggest that GFP intervention can effectively regulate the polarization imbalance of extrahepatic macrophages.

### Validation of the polarization mechanism in vitro

In order to further elucidate the underlying mechanisms by which Grifola frondosa ameliorates insulin resistance and hepatic inflammation in type 2 diabetes mellitus, we conducted in vitro validation using bone marrow-derived macrophages.The results of immunofluorescence showed that CD68 was expressed on the membrane of BMDM, and the percentage of CD68-positive cells was visually higher (**Fig. 6A**). The results of flow cytometry showed that the rate of peripheral-derived macrophages (CD68^+^CD11b^+^) was about 91.4 ± 2.1 % (**Fig. 6B**). Indicating that the extraction of BMDM was successful and could be used for the subsequent experiments.

**Fig. 6.**
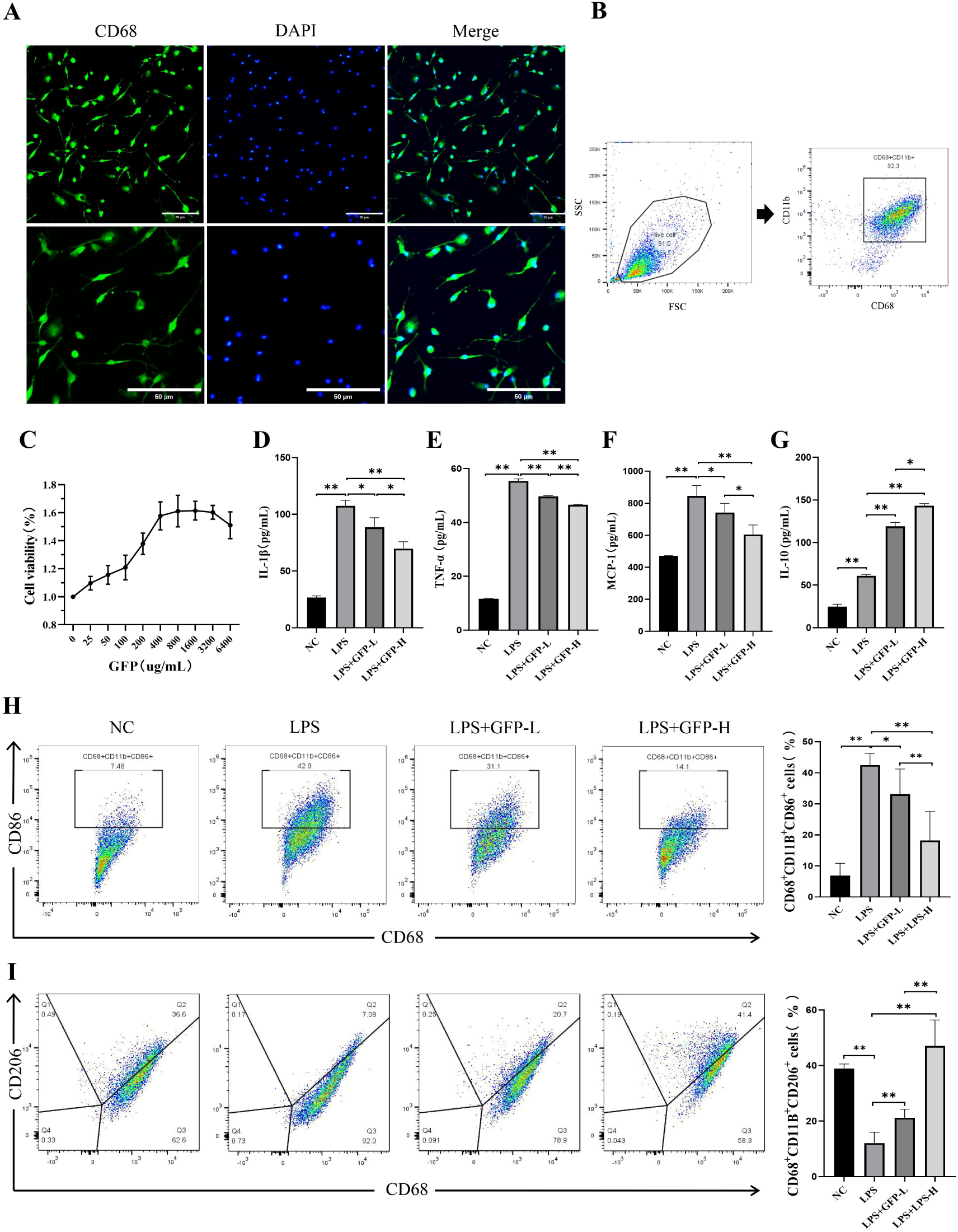
GFP inhibited M1-type polarization and promoted M2-type polarization in bone marrow-derived macrophages. (A-B) Expression of CD68 was evaluated by immunofluorescence and Flow cytometry inBMDM. (C) Effect of GFP on the proliferative activity of BMDM. (D-G) Levels of inflammatory factors in BMDM. (H) The proportion of overall M1 macrophages. (I) The proportion of overall M2 macrophages. The data were presented as mean ± SEM. * *P* > 0.05; ** *P* > 0.01; ns *P* > 0.05.

The CCK-8 results showed that GFP promoted BMDM proliferation in dose-dependent when the GFP concentration was lower than 3200 ug mL^-1^ (**Fig. 6C**). Based on the results of CCK-8 combined with previous relevant studies[21], in subsequent experiments, GFP concentrations of 200 ug mL^-1^ and 400 ug mL^-1^ were selected as intervention concentrations for the low-dose and high-dose groups, respectively.

The ELISA results showed that the levels of IL-1β, TNF-α, and MCP-1 of BMDM in the model group significantly increased, while the secretion of IL-10 was significantly decreased compared to the NC group (*P* < 0.01). After GFP intervention, the levels of IL-1β, TNF-α, and MCP-1 were significantly decreased, while the level of IL-10 was significantly increased (*P* < 0.05) in dose-dependence manner (**Fig. 6D, 6E, 6F, 6G**). It was suggested that GFP could effectively inhibit the secretion of pro-inflammatory factors and promote the secretion of anti-inflammatory factors in BMDM, which ultimately achieves the anti-inflammatory effect.

The results of flow cytometry showed that the percentage of M1-type macrophages (CD68^+^CD11b^+^CD86^+^) in the model group was significantly increased, and the rate of M2 macrophages (CD68^+^CD11b^+^CD206^+^) was significantly decreased compared to the NC group (*P* < 0.01); After GFP intervention, the percentage of M1-type macrophages was significantly decreased, and the rate of M2-type macrophage was significantly increased (*P* < 0.05) in dose-dependence manner (*P* < 0.01) (**Fig. 6H, 6I**). It suggested that GFP could inhibit M1 polarization and promote M2 polarization of BMDM.

## Discussion

T2DM is commonly characterized by abnormal blood glucose levels, weight loss, and IR; its manifestation is impaired utilization of insulin by target tissues, resulting in elevated circulating insulin levels. Its later manifestation involves the inability of pancreatic β-cells to compensate for IR, leading to pancreatic dysfunction, decreased insulin secretion, and ultimately manifested as marked hyperglycemic symptoms (Liu L et al., 2022). The high-fat diet can trigger IR in the organism. At the same time, STZ can induce pancreatic β-cell injury, and the animal model of T2DM constructed by the high-fat diet combined with STZ injection has been confirmed to have the same clinical characteristics as human diabetes (Magalhães DA et al., 2019). In this study, this approach was adopted for the construction of the T2DM rat model, and the experimental results showed that the blood glucose levels of the rats in the model group were obviously elevated and accompanied by high insulin levels and the increased insulin resistance coefficient, which was consistent with the clinical manifestations of T2DM and was an ideal animal research model for T2DM.

Typical symptoms of T2DM are persistent hyperglycemia, weight loss, and increased food and water intake (Ferguson D et al., 2021). In this study, rats in the model group showed symptoms of persistent hyperglycemia levels and weight loss compared to the NC group, this could be due to impaired energy metabolism, poor glycemic control, and a lack of carbohydrates, which causes muscle atrophy and tissue protein loss (Stanhope KL et al., 2016), GFP may improve weight loss symptoms in T2DM rats through this pathway. It is thought that insulin action is necessary to preserve glucose homeostasis. Under normal conditions, hyperinsulinemia typically coexists with hyperglycemia (Rachdaoui N et al., 2020). Besides, IPGTT and IPITT are two well-known experiments for evaluating metabolic status that can measure insulin resistance and glucose tolerance (La Valle A et al., 2023). The results of this study indicate that GFP has a positive impact on blood glucose regulation, insulin resistance improvement, and glucose tolerance restoration in T2DM rats. Specifically, GFP was observed to significantly down-regulate serum insulin levels, HOMA-IR, IPGTT-AUC, and IPITT-AUC in T2DM rats. This result is consistent with research from earlier (He X A et al., 2017).

The liver plays an important role in systemic glucose metabolism(Han HS et al., 2016). Hepatic injury caused by IR, inflammation, and oxidative stress can worsen glucose metabolism problems and accelerate the progression of T2DM (Hu W et al., 2022). The degree of liver injury can be mitigated by GFP by reducing liver inflammation, as multiple studies have demonstrated in recent years (Meng M et al., 2021). However, it has not been reported whether the aforementioned processes are related to T2DM. According to the study’s findings, the rats in the T2DM model group had a much higher liver index than the rats in the normal group, and their liver inflammation and IR had also dramatically worsened. Conversely, administration of GFP or metformin resulted in the resolution of the aforementioned symptoms, suggesting that GFP may reduce the degree of liver injury by improving liver inflammation and IR, which in turn promotes glucose metabolism balance and ameliorates symptoms associated with T2DM. Furthermore, the risk of T2DM is closely related to high levels of inflammatory cytokines (Liu Y et al., 2018). TNF-α and IL-1β are multifunctional cytokines that have been linked to inflammation, autoimmune processes, and the modulation of insulin signaling pathways to impact insulin sensitivity, among other inflammatory factors (Li T et al., 2023). In this study, we found that the secretion levels of TNF-α and IL-1β were significantly increased in the liver tissues of rats in the model group. In contrast, GFP intervention significantly down-regulated its secretion levels, which suggests that GFP can attenuate hepatic inflammation and insulin resistance in T2DM rats by down-regulating the secretion of TNF-α and IL-1β. IL-10 is a key anti-inflammatory cytokine produced during the activation of immune cell (Yin S et al., 2011). The study’s findings demonstrated that GFP can effectively reduce inflammation by increasing IL-10 release. The findings align with the research conducted by Chien RC et al. (Chien RC et al., 2016).

This study has demonstrated that GFP reduces hepatic inflammation in T2DM rats; nevertheless, further investigation is required to determine the underlying mechanisms. Previous studies have reported that the mechanism of liver inflammation is related to hepatocyte steatosis, oxidative stress, and mitochondrial dysfunction(Gaul S et al., 2021) (Ntamo Y et al., 2022) (Zhang IW et al., 2022). However, the exact mechanism underlying liver inflammation varies depending on the etiology. The role of macrophages and their polarization process in inflammation has also been studied recently. It has been found that macrophages can reduce the production of inflammatory cytokines by recognizing pathogen-associated molecular patterns (PAMPs) and damage-associated molecular patterns through pattern recognition receptors (PRRs) such as Toll-like receptors (TLRs) (Fernandes-Santos C et al., 2022). Furthermore, research has demonstrated that AMPK pathway activation might cause macrophage polarization to the anti-inflammatory M2-type, potentially improving inflammation in obese individuals (Patel VB et al., 2016). Research has demonstrated that inhibiting fatty acid production and Akt palmitoylation in macrophages may result in anti-inflammatory effects in addition to influencing macrophage polarization (An L et al., 2022). Hence, it remains unclear whether the reduction in hepatic inflammation and insulin resistance induced by GFP is linked to macrophage infiltration and polarization. The results of this study showed that 8 weeks of GFP intervention significantly inhibited macrophage infiltration into liver tissues. Additionally, phenotypic analysis of the macrophages showed that the proportion of M1-type macrophages labeled with CD86 decreased while the proportion of M2-type macrophages labeled with CD206 increased. Meanwhile, CD11b-labeled peripheral-derived macrophages were significantly increased in the liver tissues of T2DM rats. Notably, it was found that peripheral-derived macrophages predominated among pro-inflammatory M1-type hepatic macrophages in the livers of T2DM rats with obvious inflammatory lesions, while peripheral-derived macrophages were also found among anti-inflammatory M2-type macrophages following GFP intervention. The predominance of peripheral-derived macrophages suggests that peripheral-derived macrophages play a dominant role in the development of hepatic inflammation in T2DM rats, the result is consistent with the findings of Shaoying Zhang et al. (Zhang S et al., 2023). Peripheral-derived macrophages mainly include bone marrow-derived monocyte/macrophages, peritoneal macrophages, and splenic macrophages. Among them, bone marrow-derived macrophages (BMDM) are the main members of infiltrating macrophages and recurrent macrophages (Wang W et al., 2020). Thus, via in vitro studies, this study further elucidated the regulatory effect of GFP on macrophage polarization using BMDM as the experimental model. Taken together, the results suggest that the regulation of macrophage polarization homeostasis is most likely an important target for GFP to inhibit hepatic inflammation in T2DM rats.

## Conclusions

In this study, we examined the potential mechanisms behind GFP’s ameliorative effects on insulin resistance and hepatic inflammation in T2DM rats, with a focus on macrophage polarization. The research findings indicate that Grifola frondosa polysaccharide can alleviate liver inflammation by reducing macrophage infiltration, inhibiting M1-type polarization, and promoting M2-type polarization, ultimately ameliorating blood glucose levels and insulin resistance in T2DM rats, which providing a new evidence for the application of natural plants in the early prevention and treatment of T2DM.

## Acknowledgments

The authors thank the financial support of Guizhou Science and Technology Department Plan Project (Guizhou Science and Technology Foundation-ZK [2023General 315), the Guizhou Science and Technology Combined Support [2021] 134 and the Guizhou Province’s first-class discipline construction project, Public Healthand Preventive ledicine (number 2017).

## Data Availability Statement

Data are contained within the article.

## Author contributions

P.Z., S.W., X.L., P.L. conceived and designed the experiments; P.Z., L.W., Y.S., C.H., Y.P., X.L. performed the experiments; P.Z., L.W. analyzed the data; P.Z. wrote the paper; X.Q. article revision; S.W., P.L. funding acquisition. All authors have read and approved the manuscript for publication.

## Disclaimer/Publisher’s Note

The statements, opinions and data contained in all publications are solely those of the individual author(s) and contributor(s) and not of MDPI and/or the editor(s). MDPI and/or the editor(s) disclaim responsibility for any injury to people or property resulting from any ideas, methods, instructions or products referred to in the content.

